# The cell-autonomous clock of VIP receptor VPAC2 cells drives circadian behaviour

**DOI:** 10.1101/2020.08.02.232462

**Authors:** Ryan Hamnett, Johanna E. Chesham, Elizabeth S. Maywood, Michael H. Hastings

## Abstract

Circadian (∼daily) rhythms pervade mammalian behaviour. They are generated by cell-autonomous, transcriptional/translational feedback loops (TTFL), active in all tissues. This distributed clock network is co-ordinated by the principal circadian pacemaker, the hypothalamic suprachiasmatic nucleus (SCN). Its robust and accurate time-keeping arises from circuit-level interactions that bind its individual cellular clocks into a coherent time-keeper. Cells that express the neuropeptide vasoactive intestinal peptide (VIP) mediate retinal entrainment of the SCN, and in the absence of VIP, or its cognate receptor VPAC2, circadian behaviour is compromised because SCN cells cannot synchronise. The contributions to SCN pacemaking and circadian behaviour of other cell types, not least the VPAC2-expressing target cells of VIP, are, however, not understood. We therefore employed intersectional genetics to manipulate the cell-autonomous TTFL of VPAC2-expressing cells, creating temporally chimaeric mice. We could then determine whether and how VPAC2-expressing cells (a minority ∼35% of SCN cells) contribute to SCN time-keeping. Lengthening of the intrinsic TTFL period of VPAC2 cells by deletion of the *CK1ε*^*Tau*^ allele concomitantly lengthened the period of circadian behavioural rhythms. It also increased the variability of the circadian period of bioluminescent TTFL rhythms in SCN slices recorded *ex vivo*. Abrogation of circadian competence in VPAC2 cells by deletion of *Bmal1* severely disrupted circadian behavioural rhythms and compromised TTFL time-keeping in the corresponding SCN slices. Thus, VPAC2-expressing cells are a distinct, functionally powerful subset of the SCN circuit, contributing to computation of ensemble period and maintenance of circadian robustness. These findings extend our understanding of SCN circuit topology.

## Introduction

Mammalian physiology and behaviour are rhythmic and adaptively aligned with the environmental light-dark cycle by the suprachiasmatic nucleus (SCN) of the hypothalamus, which is entrained by direct retinal innervation (Reppert and Weaver, 2002). Within the 20,000 “clock” cells of the SCN, and indeed within most cells across the body, cell-autonomous circadian timekeeping is maintained by a transcriptional/translational feedback loop (TTFL) in which CLOCK:BMAL1 heterodimers activate transcription of *Period1/2* and *Cryptochrome1/2* genes, the protein products of which (PER1/2 and CRY1/2) feedback to repress their own transcription. This cycle takes approximately 24 h to complete, although the period of the SCN can be modified considerably by genetic or pharmacological manipulation (Patton et al., 2016), such as the *Tau* mutation in casein kinase 1 epsilon (*CK1ε*^*Tau*^), which shortens it to 20 hours in homozygous mice (Meng et al., 2008). More dramatically, mice lacking BMAL1 (*Bmal1*^*-/-*^), the only non-redundant TTFL component, are arrhythmic at cellular and behavioural levels (Bunger et al., 2000): effects that are reversible by transgenic rescue (McDearmon et al., 2006).

Beyond the cell-autonomous clock, SCN pacemaking is a product of circuit-level interactions that bind the circadian cycles of the individual cells. These interactions confer onto the network its essential emergent properties of robust, high-amplitude and synchronised cellular oscillations, with established ensemble phase and period (Hastings et al., 2018). Indeed, the power of intercellular coupling can partially compensate for various genetic losses within the TTFL (Liu et al., 2007; Ko et al., 2010). Although GABA is the principal neurotransmitter of the SCN, the strongest evidence for neurochemical mediation of intercellular coupling is not for GABA-ergic signalling but, rather, for a hierarchy of neuropeptides (Maywood et al., 2011). Across the SCN, discrete populations of cells are characterised by their expression of, *inter alia*, vasoactive intestinal peptide (VIP), arginine vasopressin (AVP), gastrin-releasing peptide (GRP), prokineticin2 (Prok2), and their cognate receptors (Abrahamson and Moore, 2001; Antle and Silver, 2005; Park et al., 2016). These cell-types show a highly stereotypical spatial organization within the SCN, with VIP and GRP cells in the retinorecipient “core”, and AVP and Prok2 cells in its surrounding “shell”. Mice or SCN slices deficient in intercellular communication mediated by VIP and its receptor VPAC2 (encoded by the *Vipr2* gene), display weakened rhythmicity, fewer rhythmic neurons, and damped and desynchronised cellular oscillations (Harmar et al., 2002; Colwell et al., 2003; Aton et al., 2005; Maywood et al., 2006; Ciarleglio et al., 2009). Moreover, VIP cells receive the retinal information that entrains the SCN to solar time (Abrahamson and Moore, 2001; Jones et al., 2018; Mazuski et al., 2018). In turn, VIP acts via the VPAC2-expressing cells of the SCN shell to maintain steady-state circuit-level coherence and to reset ensemble phase in response to retinal input. It achieves this via a cascade of kinase-dependent signalling (including ERK1/2 and its regulator DUSP4) and consequent regulation of a broad transcriptional network (Hamnett et al., 2019). VIP is thereby able to control both cell-autonomous and circuit-level circadian oscillations within the SCN.

The VIP/VPAC2 axis is, therefore, a central element of SCN circuit topology. Although some functions of VIP cells are established, the functions of their target cells expressing VPAC2, which constitute the next step in the SCN synaptic circuitry, are not. Located in the SCN shell, they may mediate circadian output from the SCN, and/or they may contribute to the circuit-level computations that generate its emergent properties. To investigate this, we employed transgenic mice in which VPAC2 cells express Cre recombinase (Patton et al., 2020). This allowed conditional manipulation of the cell-autonomous TTFL of VPAC2-expressing cells, altering their intrinsic period by deletion of *CK1ε*^*Tau*^, or their circadian competence by deletion of *Bmal1*. By monitoring the consequences for behaviour and SCN pacemaking, we reveal that VPAC2-expressing cells are a distinct, functionally powerful subset of the SCN circuit, contributing to computation of ensemble period and maintenance of robustness. These findings extend our understanding of SCN circuit topology.

## Materials & Methods

### Animals

All animals were cared for in accordance with the UK Animals (Scientific Procedures) Act of 1986 with local ethical approval (LMB Animal Welfare and Ethical Review Body). VPAC2-Cre mice (Tg(Vipr2cre)KE2Gsat/Mmucd; RRID:MMRRC_034281-UCD) were purchased from GENSAT (Gene Expression in the Nervous System Atlas) project (Rockefeller University, New York City, USA). These mice were subsequently crossed with either *Ck1ε*^*Tau/Tau*^ (Meng et al., 2008) or *Bmal1*^*flx/flx*^ mice (generated from Jackson Labs mouse stock no. 007668, RRID:IMSR_JAX:007668). Both the *Ck1ε*^*Tau/Tau*^ and *Bmal1*^*flx/flx*^ mice contain floxed exons that can be removed through Cre-mediated recombination. Due to VPAC2-Cre expression in developing spermatocytes (Usdin et al., 1994; Krempels et al., 1995), which is a known issue with some Cre driver lines (Luo et al., 2020), recombination occurred prior to fertilization, resulting in offspring containing one recombined allele in all cells, alongside the remaining floxed allele to be deleted only in somatic cells expressing VPAC2-Cre. These crosses, therefore, generated VPAC2-Cre/*Ck1ε*^*Tau/-*^ and VPAC2-Cre/*Bmal1*^*flx/-*^ mice. All mice were also crossed to PER2::LUCIFERASE knock-in mice (gift from Prof Joseph Takahashi (Yoo et al., 2004; University of Texas Southwestern Medical Center, Dallas)) for visualisation of circadian dynamics through bioluminescent recording, and R26R-EYFP mice (Jackson stock 006148, RRID:IMSR_JAX:006148) to report Cre-mediated recombination and determine *Bmal1* deletion efficiency (Srinivas et al., 2001). Finally, Drd1a-Cre/*Bmal1*^*Flx/-*^ were generated by crossing dopamine 1A-receptor (Drd1a)-Cre (GENSAT, RRID:MMRRC_030779-UCD) with *Bmal1*^*flx/-*^ mice. This Cre line has previously been shown to have extensive expression in the SCN, covering 63% of SCN cells and colocalising with 62% of AVP cells and 81% of VIP cells (Smyllie et al., 2016). The Drd1a-Cre population has also been shown as being capable of dictating period by crossing with *Ck1ε*^*Tau/Tau*^ mice. Given its expression profile, it served here as a comparator control for *Bmal1* deletion in VPAC2-Cre cells.

### Mouse wheel-running behaviour and analysis

Because the circadian behaviour of adult female mice is modulated by the oestrous cycle, and the oestrous cycle is itself a product of SCN circadian timekeeping, this project used only male mice to avoid potential indirect effects on behaviour arising from SCN-oestrous-behaviour interactions. Male mice were individually housed and kept in a ventilated stainless-steel cabinet with controlled lighting for the duration of behavioural monitoring. Their activity patterns were assessed using running-wheels (Actimetrics Inc.) and passive infrared movement detectors. Mice were typically entrained to a cycle of 12 hours light (L, ∼200 lx) and 12 hours dim red light (D, <10Lx) (12:12 LD) for at least 7 days to assess entrainment to 24 h rhythms, before being transferred to constant dim red light (DD) to investigate free-running period. Food and water were provided *ad libitum*. Wheel revolutions and general movement data were acquired and stored in six-minute bins. Data were analysed using ClockLab v6 (ActiMetrics Inc.; RRID:SCR_014309) with behavioural circadian period in different lighting conditions determined by Chi-squared periodogram. The emergence of the arrhythmic/disordered phenotype in *Bmal1*-deleted mice was determined by eye. The robustness of the rhythms of such mice was quantified using the Relative Amplitude (RA) non-parametric measure in ClockLab v6 on the final 14 days of DD recording.

### SCN organotypic slices: bioluminescent recordings and analysis

SCN organotypic slices and media formulations were prepared as described in Hastings et al. (2005). Briefly, brains were dissected from adult mice and placed into ice-cold dissection medium. SCN tissue was isolated from 300 µm slices prepared using a McIlwain Tissue Chopper (RRID:SCR_015798) and cultured on a Millicell filter membrane (Millipore, RRID:SCR_015799) in 1 ml culture medium. SCN slices acclimatised for 3-6 h at 37°C, 5% CO_2_, and were then transferred to 35 mm culture dishes containing 1.2 ml recording medium and sealed with glass coverslips, secured with silicon grease, for bioluminescent recordings. Slices were placed under photomultiplier tubes (PMTs; H9319–11 photon-counting head, Hamamatsu) in a light-tight incubator kept at 37°C for recording of bioluminescence. These recordings were analysed to calculate circadian period, amplitude and relative amplitude error (RAE; a measure of the rhythm robustness) using the Fast Fourier Transform – Non-Linear Least Squares (FFT-NLLS) function in the BioDare2 software (Zielinski et al., 2014). A 24 h rolling average subtraction was performed on individual traces to account for variable baselines. The first 12 h of recordings was not included in the analysis to exclude potential artefacts arising from slice preparation. To track the progression of period change in *Ck1ε*^*Tau*^ experiments, the time between successive PER2::LUCIFERASE peaks was determined.

### Immunohistochemistry and image analysis

For histological analysis, adult mice were killed rapidly by cervical dislocation and the brains dissected and immediately post-fixed in 10 ml 4% paraformaldehyde (Alfa Aesar) in 0.1 M phosphate buffer for 4-5 h at room temperature, before being cryopreserved in 20% sucrose (Fisher Chemical) in PBS at 4°C overnight. Coronal sections (40 µm) were taken using a freezing microtome (Anglia Scientific), incubated for 1 h at room temperature in 2% normal serum in PBT (PBS with 1% bovine serum albumin and 0.3% Triton X-100) then transferred to primary antisera incubation overnight at 4°C (see Table 1 for antisera details). Tissue was then incubated with appropriate secondary antibodies (1:500) for 1 h at room temperature. Sections were mounted onto slides and cover-slipped using Vectashield Hardset mounting medium with DAPI (Vector Labs, RRID:AB_2336788).

**Table 1.**
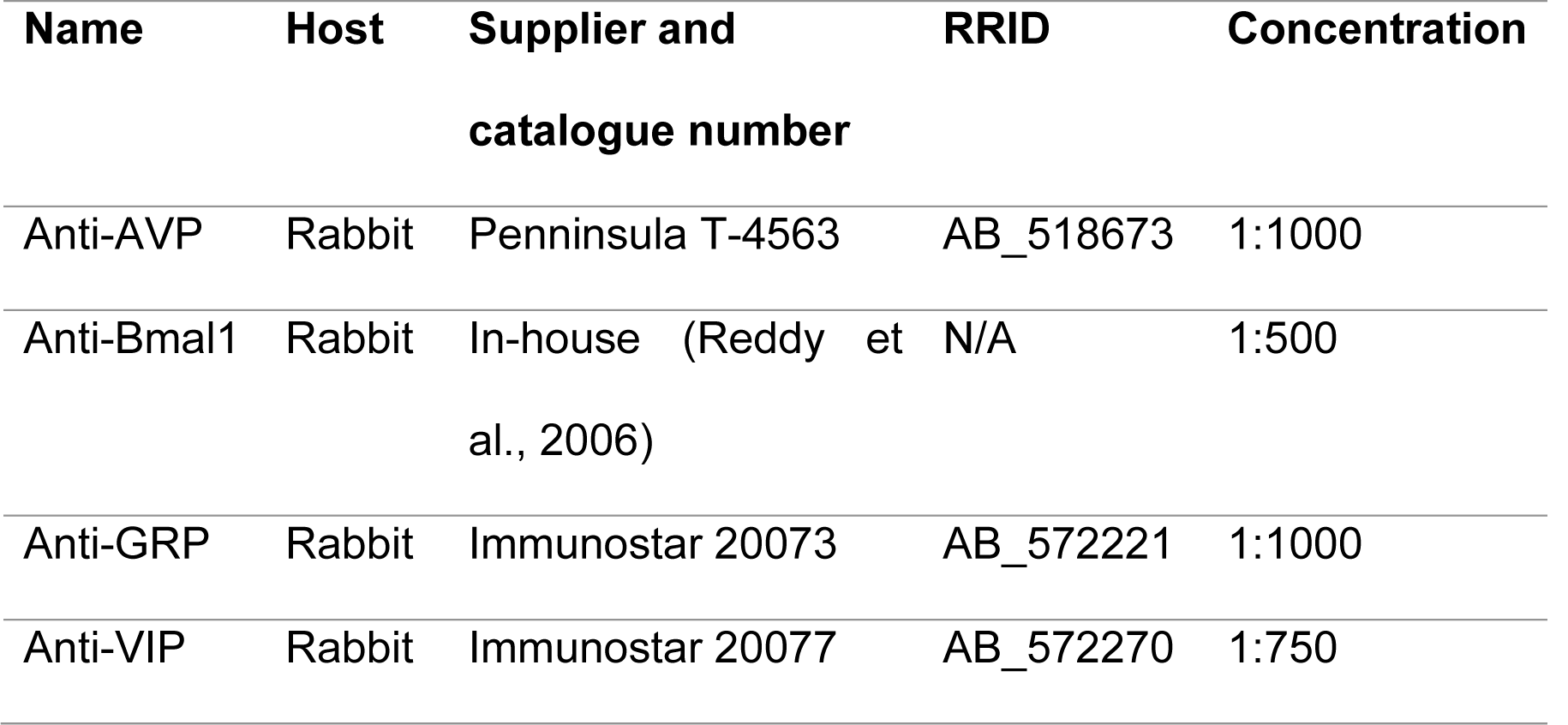
Primary antisera used for immunohistochemistry. AVP, arginine vasopressin; GRP, gastrin-releasing peptide; VIP, vasoactive intestinal peptide

Fluorescence imaging was conducted using Zeiss 710 and 780 confocal microscopes. Whole SCN sections were imaged using a 20x air objective (NA 0.5 or 0.8 on the 710 and 780 microscopes respectively) while more detailed images (required for cell-counting analysis) were acquired with a 63x oil objective, NA 1.4 before subsequent automated tile-stitching was performed by the Zeiss software (Zen 9 or 10). Counts of cells in SCN sections and determination of fluorescence intensity of SCN neuropeptides were performed in FIJI.

### Experimental design and statistical analysis

Statistical tests and graphical representation of data (mean ±SEM) were performed using Prism 6 and 7 software (GraphPad). Statistical comparisons were performed using one- or two-way ANOVA with Tukey’s multiple comparisons correction, unless otherwise stated. Kruskal-Wallis tests were performed when comparing RA scores between genotypes followed by Dunn’s correction for multiple comparisons. Correlation was determined using Pearson’s correlation coefficient. Mice and SCN slices were assigned randomly, without regard to genotype, to activity-recording cages and to photomultiplier recording systems, respectively. For both mice *in vivo* and SCN slices *ex vivo*, procedures were performed simultaneously on all genotypic groups within an experimental cohort.

## Results

### VPAC2 cells determine the period of the in vivo circadian rhythm of wheel-running behaviour

The period-setting potential of VPAC2 cells was examined by generating temporally chimaeric mice using intersectional genetics to delete the *Tau* allele of the *Ck1ε* gene in VPAC2-expressing cells. *Ck1ε*^*Tau*^ is a semi-dominant point mutation that causes a gain of function for the CK1ε protein that shortens the wild-type behavioural period by 2 h per copy (heterozygote: 22 h period; homozygote: 20 h) (Meng et al., 2008). In contrast, mice lacking *Ck1ε* (*Ck1ε*^*-/-*^), following Cre-mediated excision of a floxed exon carrying the mutation, have a 24 h period, whilst *Ck1ε*^*Tau/-*^ exhibit a 22 h period. Hence, in the VPAC2-Cre/*Ck1ε*^*Tau/-*^ mouse, VPAC2-Cre cells would have a period of 24 h (being *Ck1ε*^*-/-*^), whereas the rest of the SCN and other tissues would retain a period of 22 h (Fig. 1*A*). If the cell-autonomous TTFL of VPAC2 cells directs circadian behaviour, such lengthening of their intrinsic period should correspondingly lengthen the period of wheel-running rhythms of the mouse.

**Figure 1.**
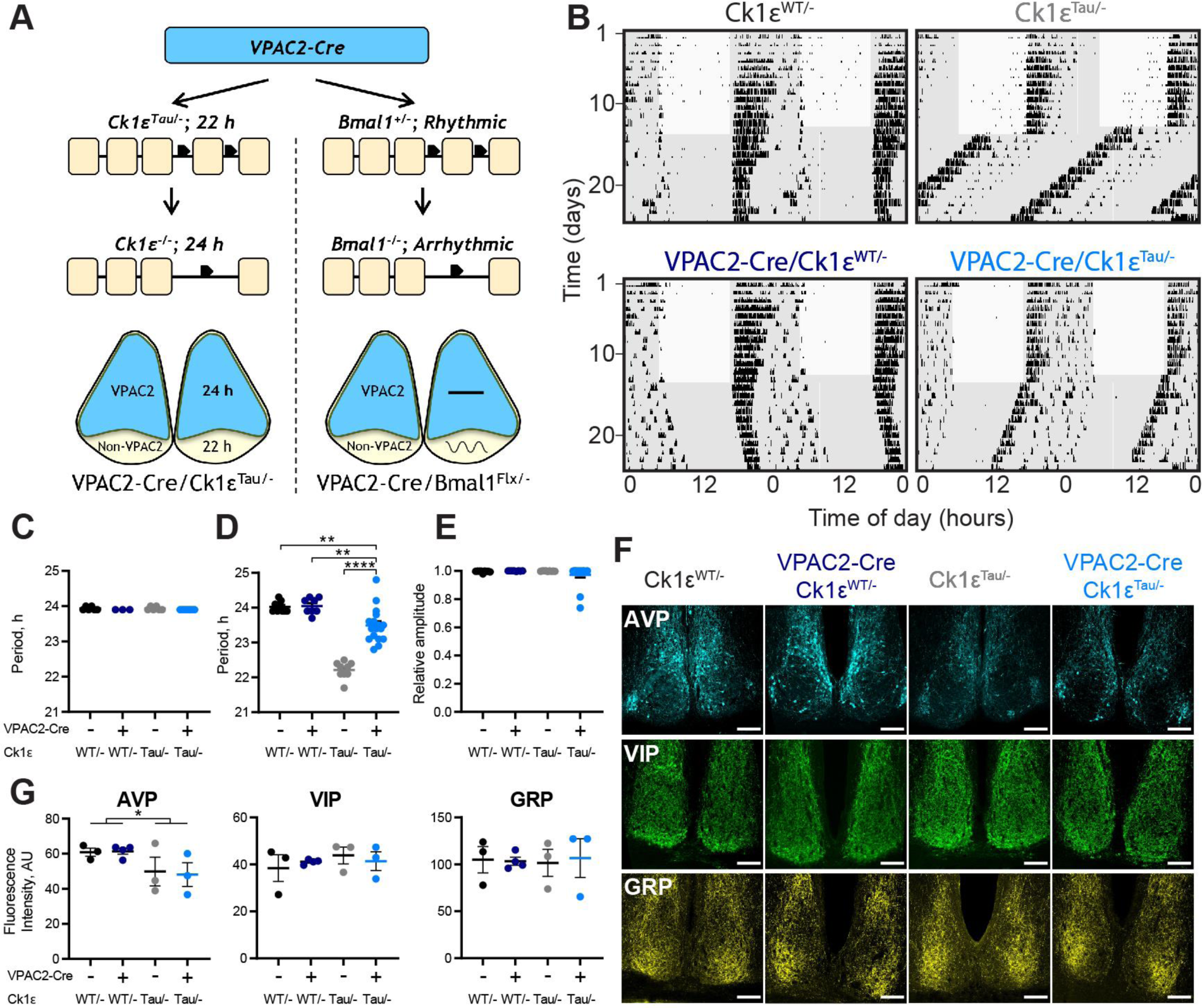
Intersectional genetics reveals that VPAC2-expressing cells determine the circadian period of mouse wheel-running behaviour. ***A***, Schematic view of the intersectional approach, whereby mice carrying Cre recombinase as a transgene controlled by the VPAC2 promoter were crossed with either *Ck1ε*^*Tau/-*^ or *Bmal1*^*flx/-*^ mice to create a chimaeric SCN. The result is for VPAC2 cells to lack either the Tau allele or Bmal1, reverting them to a 24 h period as *Ck1ε*^*-/-*^ or rendering them arrhythmic as *Bmal1*^*-/-*^, respectively. ***B***, Representative double-plotted actograms of wheel-running activity of VPAC2-Cre/*Ck1ε*^*Tau/-*^ mice and their respective controls exposed to a 12:12 light:dim red (LD) cycle followed by continuous dim red light (DD) conditions. Grey shading represents dim red. ***C, D*** Periods (mean ±SEM) observed in 12:12 LD (***C***; n = 6 (*Ck1ε*^*WT/-*^), 3 (VPAC2-Cre/*Ck1ε*^*WT/-*^), 6 (*Ck1ε*^*Tau/-*^) and 7 (VPAC2-Cre/*Ck1ε*^*Tau/-*^)) and DD (***D***; n = 9, 8, 9, 17) of VPAC2-Cre/*Ck1ε*^*Tau/-*^ mice and their respective controls. ***E***, Relative amplitude scores (mean ±SEM) from 10 days of wheel running activity under DD (n as in ***D***). Kruskal-Wallis test. ***F***, Representative images of immunohistochemical staining of SCN of VPAC2-Cre/*Ck1ε*^*Tau/-*^ mice and their respective controls for AVP-ir (top), VIP-ir (middle) and GRP-ir (bottom). Scale bar represents 100 µm. ***G***, Fluorescence intensities (mean ±SEM) for immunohistochemical staining of AVP-ir (left), VIP-ir (middle) and GRP-ir (right) in SCN sections of VPAC2-Cre/*Ck1ε*^*Tau/-*^ mice with their respective controls (n = 3 per genotype). All tests were two-way ANOVAs (main effects of Cre and *Tau* allele) with Tukey’s multiple comparisons test, \**p* < 0.05, \*\**p* < 0.01, \*\*\*\**p* < 0.0001.

The wheel-running behaviour of mice of all genotypes (*Ck1ε*^*WT/-*^ and *Ck1ε*^*Tau/-*^, without and with VPAC2-Cre) entrained stably to a 12:12 light:dim red light (LD) cycle (Fig. 1*B, C*; two-way ANOVA, *F*_(1,18)_ = 3.706, *p* = 0.07 (Cre), *F*_(1,18)_ = 0.000, *p* > 0.999 (Tau), *F*_(1,18)_ = 0.000, *p* > 0.999 (interaction)). On transfer to continuous dim red light (DD), *Ck1ε*^*WT/-*^ mice free-ran with a wild-type equivalent period (∼24 h), whereas that of *Ck1ε*^*Tau/-*^ mice was significantly shorter (∼22 h, Fig. 1*B,D*; two-way ANOVA, *F*_(1,39)_ = 30.46, *p* < 0.0001 (interaction), Tukey *post-hoc p* < 0.0001). The presence of VPAC2-Cre had no effect in *Ck1ε*^*WT/-*^ mice lacking the floxed allele, whereas its presence in the *Ck1ε*^*Tau/-*^ mice caused significant lengthening of circadian period (Tukey *post-hoc Ck1ε*^*Tau/-*^ (22.21 ±0.08 h) vs. VPAC2-Cre/*Ck1ε*^*Tau/-*^ (23.49 ±0.12 h) *p* < 0.0001). This VPAC2-Cre/*Ck1ε*^*Tau/-*^ group mean was slightly, albeit significantly, below the 24 h period of *Ck1ε*^*WT/-*^ controls, with or without VPAC2-Cre (Tukey *post-hoc Ck1ε*^*WT/-*^ vs. VPAC2-Cre/*Ck1ε*^*Tau/-*^ *p* = 0.004). Nevertheless, at a behavioural level, the cell-autonomous clock of VPAC2 cells can exert a clear influence on circadian period, lengthening it in line with their intrinsic period. Furthermore, the coherence and amplitude of the behavioural rhythm were not affected by temporal chimaerism (Fig. 1*E*; Kruskal-Wallis test, H(3) = 3.661, *p* = 0.30), confirming that circuit-level mechanisms are able to sustain coherent circadian output in the face of widely divergent cell-autonomous periods in the SCN (Smyllie et al., 2016; Brancaccio et al., 2017).

To test further the effect of chimaerism on SCN integrity, sections of adult mouse brain were processed for immunohistochemical (IHC) staining of the neuropeptides VIP, AVP and GRP. The morphology of the SCN was comparable across all genotypes and there were no significant differences in the expression of VIP (two-way ANOVA, *F*_(1,9)_ = 0.001, *p* = 0.97 (Cre), *F*_(1,9)_ = 0.628, *p* = 0.45 (Tau), *F*_(1,9)_ = 0.518, *p* = 0.49 (interaction)) or GRP (Fig. 1*F,G*; two-way ANOVA, *F*_(1,9)_ = 0.016, *p* = 0.90 (Cre), *F*_(1,9)_ = 0.000, *p* = 0.99 (Tau), *F*_(1,9)_ = 0.062, *p* = 0.81 (interaction)). Compared to *Ck1ε*^*WT/-*^ controls, however, the *Ck1ε*^*Tau/-*^ mice showed a small (∼20%) but significant reduction of AVP expression (Fig. 1*F,G*; two-way ANOVA, *F*_(1,9)_ = 0.013, *p* = 0.91 (Cre), *F*_(1,9)_ = 5.751, *p* = 0.04 (Tau), *F*_(1,9)_ = 0.053, *p* = 0.82 (interaction)). This was evident in SCN from mice both without and with the VPAC2-Cre and so could not be a direct cause of the period-lengthening observed in the latter. Temporal chimaerism did not, therefore, affect the structural integrity of the SCN, nor the quality of behavioural circadian output, and revealed the pace-setting ability of SCN VPAC2 cells.

### Period determination by VPAC2 cells is attenuated in the SCN ex vivo

Following the recording of wheel-running behaviour, mice were killed under dim red light and their SCN PER2-driven bioluminescence rhythms were recorded to determine the impact of temporal chimaerism on the intrinsic SCN TTFL. SCN of all four genotypes exhibited robust and coherent bioluminescent rhythms (Fig. 2*A*). As with behavioural periods, the rhythms were shortened by approximately 2 h by the presence of a single copy of the *Tau* allele in the SCN of mice lacking VPAC2-Cre (Fig. 2*A,B*; two-way ANOVA, *F*_(1,21)_ = 39.01, *p* < 0.0001 (Tau)). In contrast to the lengthening of period observed *in vivo*, however, there was no systematic or significant effect of VPAC2-Cre on the mean circadian period of *Ck1ε*^*Tau/-*^ SCN slices (two-way ANOVA, *F*_(1,21)_ = 0.408, *p* = 0.53 (interaction)), and the mean circadian period was not statistically different from *Ck1ε*^*Tau/-*^ slices lacking VPAC2-Cre. There was, however, a marked variability within the VPAC2-Cre/*Ck1ε*^*Tau/-*^ group, with a range of ∼6.3 hours, a clear departure from the behavioural data (Fig. 2*B,C*). Furthermore, in the control groups there was a direct correspondence between the circadian periods measured *in vivo* as behaviour and *ex vivo* as SCN bioluminescence within individual mice, with slices typically having a slightly longer period (Fig. 2*C*). This was not the case for the VPAC2-Cre/*Ck1ε*^*Tau/-*^ group, in which the *ex vivo* SCN periods were not only widely divergent, but were also not consistent with the *in vivo* period of the corresponding mouse, typically being shorter (Fig. 2*C*; two-way ANOVA, *F*_(3,19)_ = 8.975, *p* = 0.0007 (interaction); Sidak’s post-hoc *p* < 0.0001 (VPAC2-Cre/*Ck1ε*^*Tau/-*^, Behaviour vs. Slice period)), and comparable to or even faster than those of *Ck1ε*^*Tau/-*^ SCN lacking recombinase. Consequently, when *in vivo* and *ex vivo* periods were plotted together, the control groups exhibited a highly significant within-animal correlation but the VPAC2-Cre/*Ck1ε*^*Tau/-*^ group did not (Fig. 2*D*; Pearson correlation, Controls: *r* = 0.923, *p* < 0.0001, VPAC2-Cre/*Ck1ε*^*Tau/-*^: *r* = 0.598, *p* = 0.07).

**Figure 2.**
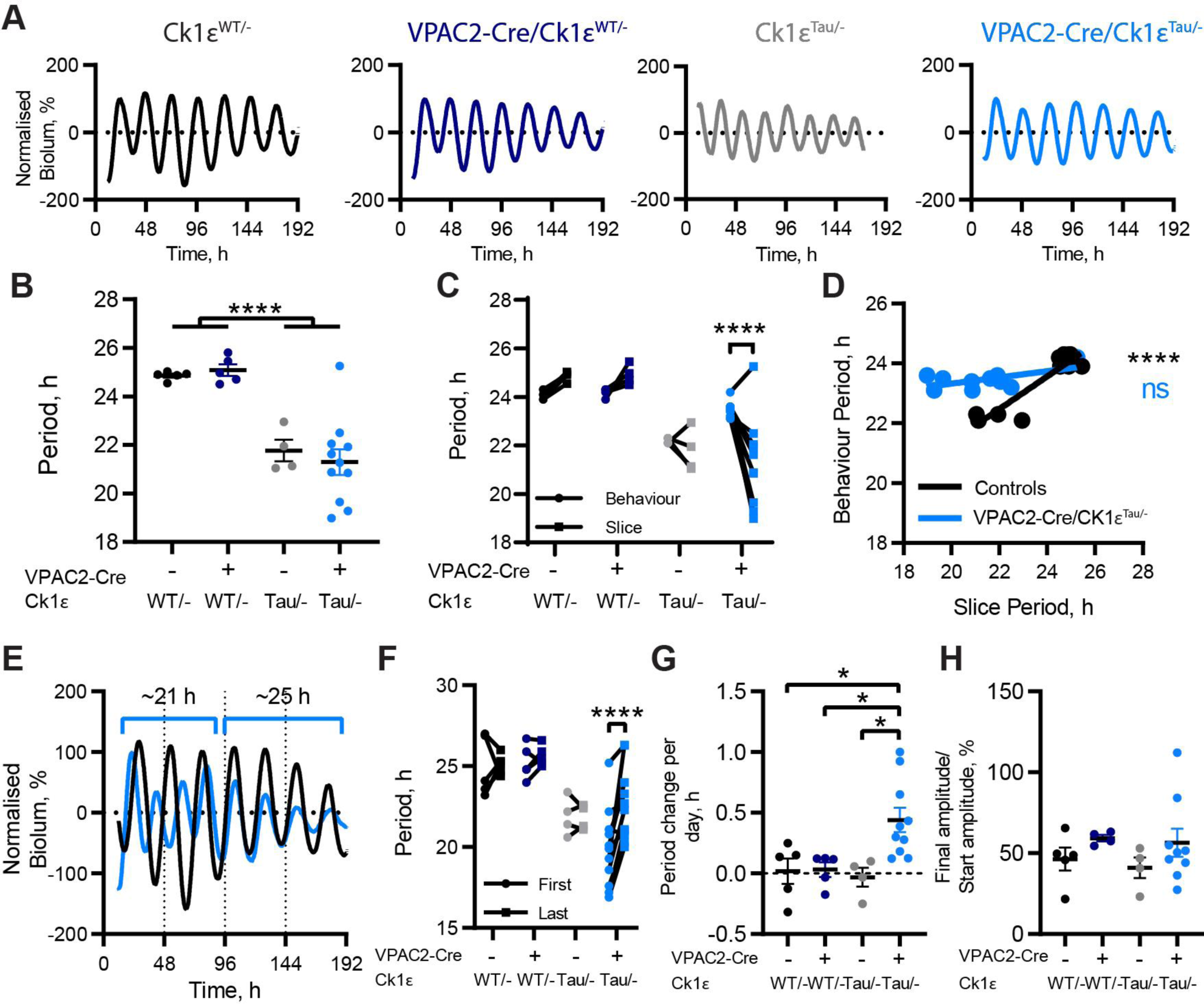
Circadian periods of bioluminescent rhythms of SCN slices from VPAC2-Cre-Ck1ε^Tau/-^ mice do not reflect behavioural period. ***A***, Representative baseline corrected PER2::LUCIFERASE bioluminescence rhythms from *Ck1ε*^*WT/-*^, VPAC2-Cre/*Ck1ε*^*WT/-*^, *Ck1ε*^*Tau/-*^ and VPAC2-Cre/*Ck1ε*^*Tau/-*^ SCN dissected following wheel-running recordings. ***B***, Periods (mean ±SEM) of the first 4 bioluminescent cycles of adult SCN slices as in ***A. C***, Comparison of circadian periods of free-running behaviour and slice bioluminescence rhythms (measured from the first 4 cycles) from individual mice. ***D***, Scatter plot of slice periods (measured from the first 4 cycles) vs. behavioural periods. VPAC2-Cre/*Ck1ε*^*Tau/-*^ mice show no significant correlation (*p* = 0.07, Pearson’s correlation) but grouped control slices do (\*\*\*\**p* < 0.0001, Pearson’s correlation). Lines represent linear regression: VPAC2-Cre/*Ck1ε*^*Tau/-*^ r^2^ =0.36, Y = 0.1037*X + 21.25; Controls r^2^ =0.85, Y = 0.5458*X + 10.47. ***E***, Representative baseline corrected PER2::LUCIFERASE bioluminescence rhythms from *Ck1ε*^*WT/-*^ and VPAC2-Cre/*Ck1ε*^*Tau/-*^ SCN slices displaying no phase alignment initially, followed by period lengthening in the VPAC2-Cre/*Ck1ε*^*Tau/-*^ slice and resultant phase alignment. ***F***, Comparison of periods between the first two bioluminescent cycles (‘First’) and final two cycles (‘Last’) within each SCN slice. ***G***, Period change per day (mean ±SEM) in SCN slices. ***H***, Bioluminescence amplitude (mean ±SEM) in the last 3 cycles as a percentage of the amplitude in the first 3 cycles. For ***B*** and ***F-H***, n = 5 (*Ck1ε*^*WT/-*^), 5 (VPAC2-Cre/*Ck1ε*^*WT/-*^), 4 (*Ck1ε*^*Tau/-*^) and 11 (VPAC2-Cre/*Ck1ε*^*Tau/-*^). For ***C*** and ***D***, n = 5 (*Ck1ε*^*WT/-*^), 4 (VPAC2-Cre/*Ck1ε*^*WT/-*^), 4 (*Ck1ε*^*Tau/-*^) and 10 (VPAC2-Cre/*Ck1ε*^*Tau/-*^). All tests were two-way ANOVAs with Tukey’s multiple comparisons test, \**p* < 0.05, \*\*\*\**p* < 0.0001.

The period-lengthening by deletion of *Ck1ε*^*Tau*^ in VPAC2 cells was therefore consistently effective *in vivo* but had variable penetrance in the corresponding SCN slices *ex vivo*. This variability was also evident in the stability of the individual SCN rhythms, whereby the period of VPAC2-*Cre/Ck1ε*^*Tau/-*^ slices tended to increase over time in culture (Fig. 2*E-G*; two-way ANOVA, *F*_(3,21)_ = 4.478, *p* = 0.014 (interaction), Tukey *post-hoc p* < 0.0001), suggesting that the long-period VPAC2 cells were re-exerting influence over ensemble period. While there was considerable variability in the peak-to-peak period within the group (Fig. 2*C,F*), the average period increase was almost 30 minutes per day for VPAC2-Cre/*Ck1ε*^*Tau/-*^ slices, whereas control slices did not show significant period-lengthening during *ex vivo* culture (Fig. 2*G*; two-way ANOVA, *F*_(1,20)_ = 4.445, *p* = 0.048 (interaction), Tukey *post-hoc p* = 0.030 (vs. *Ck1ε*^*WT/-*^), *p* = 0.023 (vs. *Ck1ε*^*Tau/-*^), *p* = 0.035 (vs. VPAC2-Cre)). Importantly, there was no significant difference in the progressive fall in amplitude observed over time in all genotypes (Fig. 2*H;* two-way ANOVA, *F*_(1,18)_ = 2.555, *p* = 0.13 (Cre), *F*_(1,18)_ = 0.216, *p* = 0.65 (Tau), *F*_(1,18)_ = 0.020, *p* = 0.89 (interaction)), suggesting that the period lengthening was not the result of cells within the slice desynchronising. Overall, we conclude that cell-autonomous properties of VPAC2 cells contribute strongly to computation of ensemble period *in vivo*, and also, but to a lesser extent, *ex vivo*.

### The cell-autonomous clock of VPAC2 cells is essential for the circadian coordination of rest/activity rhythms

To test whether circadian competence in VPAC2 cells is essential for the generation of behavioural rhythms and/or molecular pacemaking in the SCN, we examined the impact of cell-type-specific deletion of BMAL1 by Cre-mediated removal of a critical exon within a floxed allele of the *Bmal1* gene (Fig. 1*A*). Homozygous loss of *Bmal1* results in a severely disrupted or arrhythmic TTFL, although a single floxed copy of *Bmal1* is fully functional and sufficient for normal pacemaking (Bunger et al., 2000; Gibbs et al., 2012). All experiments were conducted using *Bmal1*^*flx/-*^ mice because the efficiency of *Bmal1* deletion in the SCN is enhanced when the floxed *Bmal1* allele is paired against a null allele (Husse et al., 2011). The efficiency of deletion in VPAC2-expressing cells (constituting ∼35% of SCN cells) was determined by IHC for BMAL1-expression and the expression of a genomic EYFP reporter (Rosa-LSL-EYFP) to identify Cre-expressing cells (Fig. 3*A*). VPAC2-Cre-mediated excision reduced the number of BMAL1-expressing cells across the SCN by ∼20% when compared to wild-type SCN (Fig. 3*B*; one-way ANOVA, *F*_(4,22)_ = 98.83, *p* < 0.0001, Tukey’s *post-hoc p* < 0.0001), which suggests appreciable but incomplete deletion from all VPAC2-expressing cells. Indeed, within the VPAC2-Cre expressing cell population (labelled by EYFP), ∼30% of cells had detectable BMAL1-ir. This nevertheless emphasised the efficient and specific targetting of BMAL1 in ∼70% of VPAC2 cells (Fig. 3*C*). Importantly, the loss of BMAL1 from VPAC2-expressing cells did not affect overall SCN morphology or expression of VIP or GRP (Fig. 3*E*). There was, however, a small (∼30%) but significant decline in AVP-ir (Fig. 3*D,E;* two-way ANOVA, *F*_(1, 9)_ = 14.34, *p* = 0.004 (interaction), Sidak’s *post-hoc p* = 0.004 (vs. *Bmal1*^*Flx/-*^), *p* = 0.002 (vs. VPAC2Cre/*Bmal1*^*WT/-*^)), consistent with some cells co-expressing AVP and VPAC2 (Patton et al., 2020; Wen et al., 2020), and the *Avp* gene being a circadian clock-controlled target of the BMAL1-dependent TTFL (Jin et al., 1999; Mieda et al., 2015).

**Figure 3.**
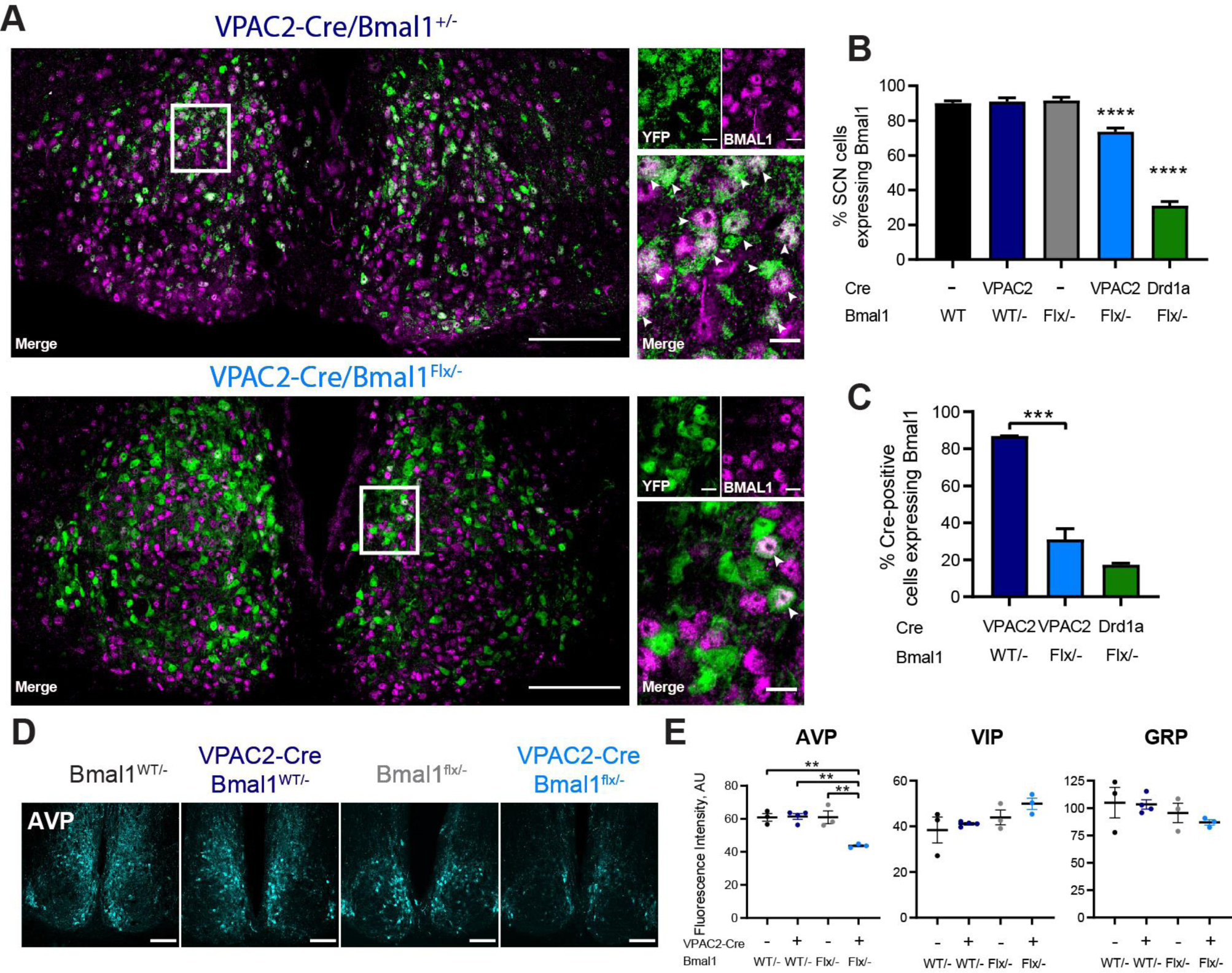
Targeted deletion of BMAL1 from VPAC2-Cre-expressing SCN cells. ***A***, Representative 63x tiled confocal micrographs of Cre recombinase activity, as reported by a genetically encoded EYFP reporter (green), and BMAL1 immunohistochemistry (magenta) in VPAC2-Cre/*Bmal1*^*WT/-*^ and VPAC2-Cre/*Bmal1*^*flx/-*^ SCN sections. Locations of magnified images are indicated by white rectangles. White arrowheads indicate co-localisation between EYFP and BMAL1-ir. Scale bars represent 100 µm in stitched images, 10 µm in magnified images. ***B***, Percentage of SCN cells (marked by DAPI; mean ±SEM) expressing BMAL1-ir across genotypes, including Drd1a-Cre/*Bmal1*^*flx/-*^. ***C***, Percentage of Cre-positive cells (marked by EYFP; mean ±SEM) expressing BMAL1-ir across genotypes. (***B, C***; One-way ANOVA with Tukey’s multiple comparisons test). ***D***, Representative images of AVP-ir in SCN of VPAC2-Cre/*Bmal1*^*flx/-*^ mice and controls. Scale bar represents 100 µm. ***E***, Fluorescence intensities (mean ±SEM) for immunohistochemical staining of AVP (left), VIP (middle) and GRP (right) in SCN sections of VPAC2-Cre/*Bmal1*^*flx/-*^ mice and controls. For ***B*** and ***C***, n = 7 (*Bmal1*^*WT/-*^), 7 (*Bmal1*^*flx/-*^), 2 (VPAC2-Cre/*Bmal1*^*WT/-*^), 8 (VPAC2-Cre/*Bmal1*^*flx/-*^) and 3 (Drd1a-Cre/*Bmal1*^*flx/-*^). For ***E***, n = 3 per group. Two-way ANOVA with Tukey’s post-hoc test. \*\**p* < 0.01, \*\*\**p* < 0.001, \*\*\*\**p* < 0.0001.

Mice of all genotypes: *Bmal1*^*WT/-*^ and *Bmal1*^*Flx/-*^ both without and with VPAC2-Cre, entrained stably under a 12:12 LD cycle with a period of 24 h and there were no obvious differences in activity patterns between them (Fig. 4*A;* two-way ANOVA, *F*_(1,29)_ = 0.459, *p* = 0.5 (Cre), *F*_(1,29)_ = 0.672, *p* = 0.42 (Bmal1), *F*_(1,29)_ = 2.81, *p* = 0.10 (interaction)). Mice were then transferred to DD for up to 7 weeks to allow the emergence of any *Bmal1*^*-/-*^-dependent phenotypes. Control mice (*Bmal1*^*WT/-*^ with or without VPAC2-Cre, and *Bmal1*^*Flx/-*^ without Cre) free-ran with clear circadian patterns and endogenous periods slightly longer than 24 h (Fig. 4*B,C*) and with well-defined amplitude (Fig. 4*E*). The VPAC2-Cre*/Bmal1*^*flx/-*^ group, however, displayed highly variable phenotypes both between and within individual animals under DD (Fig. 4*C*). Thirteen of 18 mice showed a strong phenotype: arrhythmicity occurred in 5 mice, while fragmented or “split” behaviour was the most common result, seen in 8 mice, with several animals displaying multiple significant periods (Fig. 4*B-D*). Furthermore, in the majority of mice, the amplitude of the activity rhythm was reduced, leading to a significant difference with control groups (Fig. 4*E*; Kruskal-Wallis test, H(3) = 21.19, *p* < 0.0001, Dunn’s *post hoc p* < 0.0001 (vs. VPAC2-Cre/*Bmal1*^*WT/-*^), *p* = 0.032 (vs. *Bmal1*^*Flx/-*^)). Strikingly, BMAL1-dependent phenotypes (arrhythmicity, multiple periodicities, loss of amplitude) did not appear at a consistent point following transfer to DD. Rather, they accumulated progressively (Fig. 4*F*), at a range of times throughout the experiment: some mice immediately showed split or arrhythmic behaviour, while others took several weeks and displayed phenotypes that looked wild-type until that point. We conclude that the cell-autonomous rhythmicity of VPAC2-expressing cells is essential for normal circadian control of behaviour, though considerable plasticity exists within and between individual mice that affects the precise timing and presentation of the *VPAC2-Cre/Bmal1*^*flx/-*^ phenotype.

**Figure 4.**
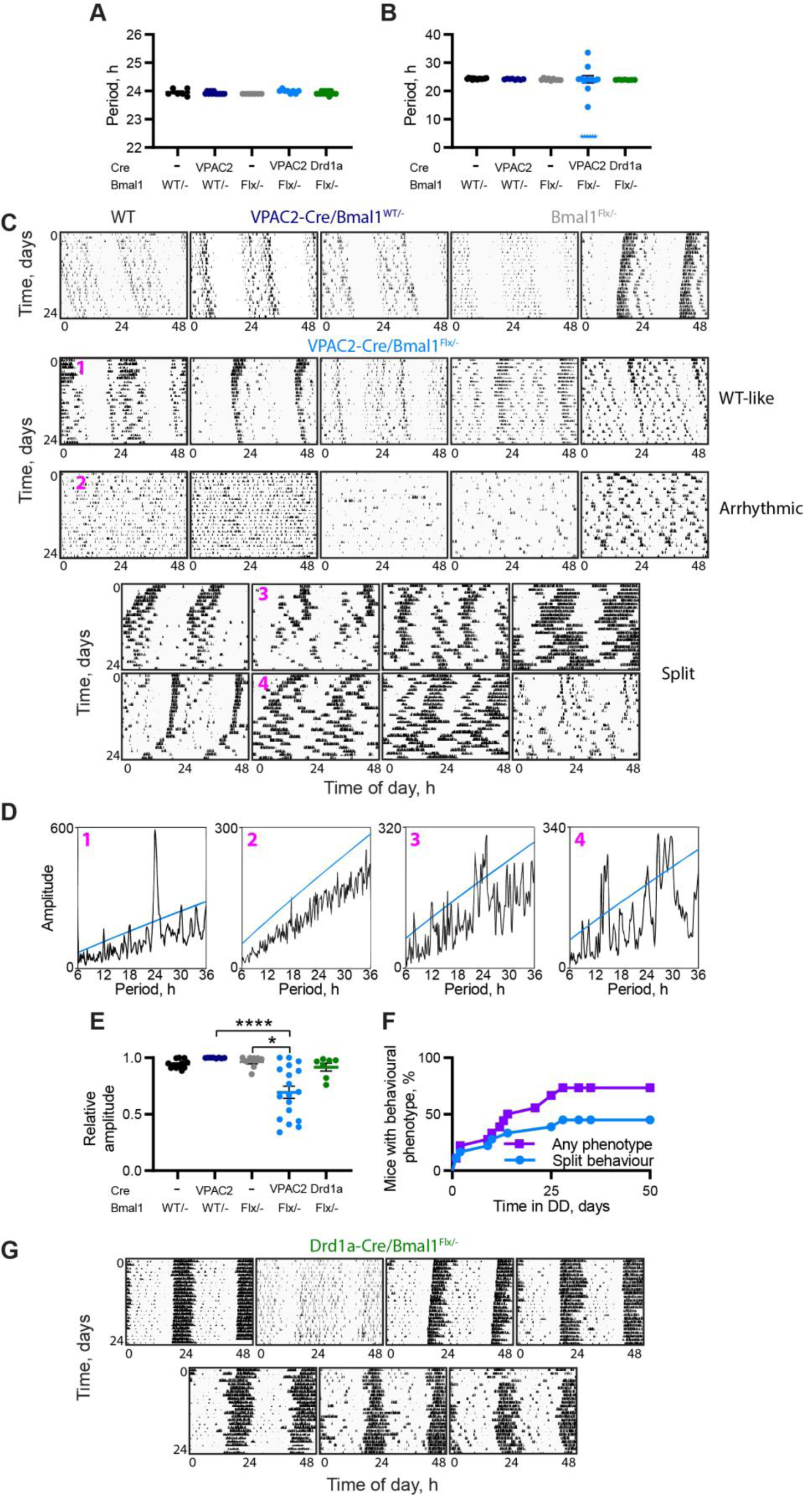
Deletion of BMAL1 from VPAC2-Cre-expressing cells compromises circadian behaviour. ***A, B***, Circadian periods (mean ±SEM) of VPAC2-Cre/*Bmal1*^*flx/-*^ mice and controls under (***A***) 12:12 LD and (***B***) DD. For 6 VPAC2-Cre/*Bmal1*^*flx/-*^ mice an explicit single period could not be determined due to their disturbed behaviour and so these are added as star symbols as a nominal 3 hours, and were excluded from statistical analysis. Two-way ANOVA (excluding Drd1a-Cre/*Bmal1*^*flx/-*^ mice); one-way ANOVA including all groups for Drd1a-Cre/*Bmal1*^*flx/-*^ comparison. ***C***, Representative double-plotted actograms of final 24 days of wheel-running activity of control mice (1 *Bmal1*^*WT/-*^, 2 VPAC2-Cre/*Bmal1*^*WT/-*^, and 2 *Bmal1*^*flx/-*^) and of all VPAC2-Cre/*Bmal1*^*flx/-*^ mice. VPAC2-Cre/*Bmal1*^*flx/-*^ actograms are divided into three groupings (WT-like, Arrhythmic, and Split) and then ranked in descending order of relative amplitude score under DD (final 14 days of recording). ***D***, Chi-square periodograms for 4 VPAC2-Cre/*Bmal1*^*flx/-*^ as indicated in ***C***. (1) has a single significant peak at ∼24 h, (2) has no significant period, and (3) and (4) display multiple significant periods. ***E***, Relative amplitude scores (mean ±SEM) from the last 14 days of wheel running activity under DD. Kruskal-Wallis test with Dunn’s multiple comparisons. n = 12 (*Bmal1*^*WT/-*^), 8 (*Bmal1*^*flx/-*^), 8 (VPAC2-Cre/*Bmal1*^*WT/-*^), 18 (VPAC2-Cre/*Bmal1*^*flx/-*^) and 7 (Drd1a-Cre/*Bmal1*^*flx/-*^). ***F***, Percentage of VPAC2-Cre/*Bmal1*^*flx/-*^ mice displaying behavioural phenotypes (purple, squares) or specifically split behaviour (blue, circles) over time. ***G***, Double-plotted actograms of wheel-running activity of all Drd1a-Cre/*Bmal1*^*flx/-*^ mice. \**p* < 0.05, \*\*\*\**p* < 0.0001.

To determine whether the behavioural effects of the local loss of BMAL1 in VPAC2 cells were specific to that cell type, an additional group of mice was included in which BMAL1 was deleted from cells expressing Cre driven by the Drd1a promoter. In these animals, BMAL1 was deleted in ∼80% of Drd1a-specific cells compared to ∼70% of VPAC2-expressing cells in VPAC2-Cre/*Bmal1*^*Flx/-*^ mice (Fig. 3*C*), resulting in an overall deletion of BMAL1 across ∼70% of total SCN cells in Drd1a-Cre/*Bmal1*^*Flx/*^ mice (Fig. 3*B*; one-way ANOVA, *F*_(4,22)_ = 98.83, *p* < 0.0001, Tukey’s *post-hoc p* < 0.0001 (vs. *Bmal1*^_*WT/-*_^ or VPAC2-Cre/*Bmal1*^*Flx/-*^)). Notwithstanding this broader deletion of BMAL1, the rest/activity rhythms of these mice were comparable to those of the control groups: only 3 of 7 showed minor instability in the time of onset of wheel-running, but none of them showed a phenotype comparable to those of VPAC2-Cre/*Bmal1*^*Flx/-*^ mice (Fig. 4*G*), and the relative amplitudes of their rhythms were not significantly different from control groups (Fig. 4*E*; Dunn’s *post hoc p* > 0.9999 (vs. *Bmal1*^*Flx/-*^)). These results emphasize that the effects of deletion of BMAL1 from VPAC2 cells are not merely due to reaching a threshold number of SCN neurons, given that the Drd1a-Cre-mediated removal targeted over twice as many cells as that of VPAC2-Cre. Rather, they suggest that the nature and identity of the cells from which BMAL1 is deleted (in this case VPAC2 or Drd1a) are paramount.

### The cell-autonomous clock of VPAC2 cells is essential for molecular pacemaking in the SCN

The loss of behavioural rhythms in conditionally BMAL1-deleted mice could indicate that a functional cell-autonomous TTFL in VPAC2 cells is required for either producing a coherent ensemble signal within the SCN circuit or for enabling distribution of an appropriate output signal from the SCN to relevant brain centres. To test this, SCN slices were prepared from adult mice following the recording of wheel-running rhythms to establish if behavioural phenotypes were reflected by, and thus likely resulted from, VPAC2-mediated changes in SCN rhythmicity. Circadian rhythms of PER2-Luc bioluminescence from slices lacking either the floxed *Bmal1* allele or VPAC2-Cre were stable and of high amplitude (Fig. 5*A,B*). In contrast, VPAC2-Cre/*Bmal1*^*Flx/-*^ SCN slices were highly disorganised, showing erratic and unstable bioluminescence (Fig. 5*A*), and having significantly higher relative amplitude error (RAE) scores compared to controls (Fig. 5*C*; one-way ANOVA, *F*_(4,32)_ = 8.147, *p* = 0.0001, Tukey’s *post-hoc p* = 0.005 (vs. *Bmal1*^*Flx/-*^), *p* = 0.0004 (vs. VPAC2Cre/*Bmal1*^*WT/-*^)). The circadian behaviour of VPAC2-Cre/*Bmal1*^*Flx/-*^ mice was therefore directly reflected in the lack of competence of molecular pacemaking of the TTFL SCN rhythm: a disrupted, unstable behavioural rhythm was predictive of a poorly organised rhythm in the SCN from the same animal (Fig. *5D*). In contrast to the disruption following deletion of BMAL1 from VPAC2 cells, but consistent with minor behavioural effects, the more extensive Drd1a-Cre-mediated deletion of BMAL1 had little impact on the SCN rhythms. They were well defined (Fig. 5*A*) with a period (Fig. 5*B*; one-way ANOVA, *F*_(4,32)_ = 1.425, *p* = 0.93) and a low RAE score comparable to those of the control groups (Fig. 5*C*; Tukey’s *post-hoc p* = 0.77 (vs. *Bmal1*^*Flx/-*^), *p* = 0.54 (vs. *Bmal1*^*WT/-*^)). The competence of their SCN was therefore consistent with their well organised wheel-running behaviour. These results confirm the specificity of the contribution of VPAC2-expressing cells to the ensemble rhythm and circadian behaviour, whereby the ablation of the cell-autonomous clock of VPAC2 cells is sufficient to abrogate SCN, and thence behavioural, rhythmicity. This highlights VPAC2-expressing cells as necessary determinants of SCN circadian output and behavioural regulation.

**Figure 5.**
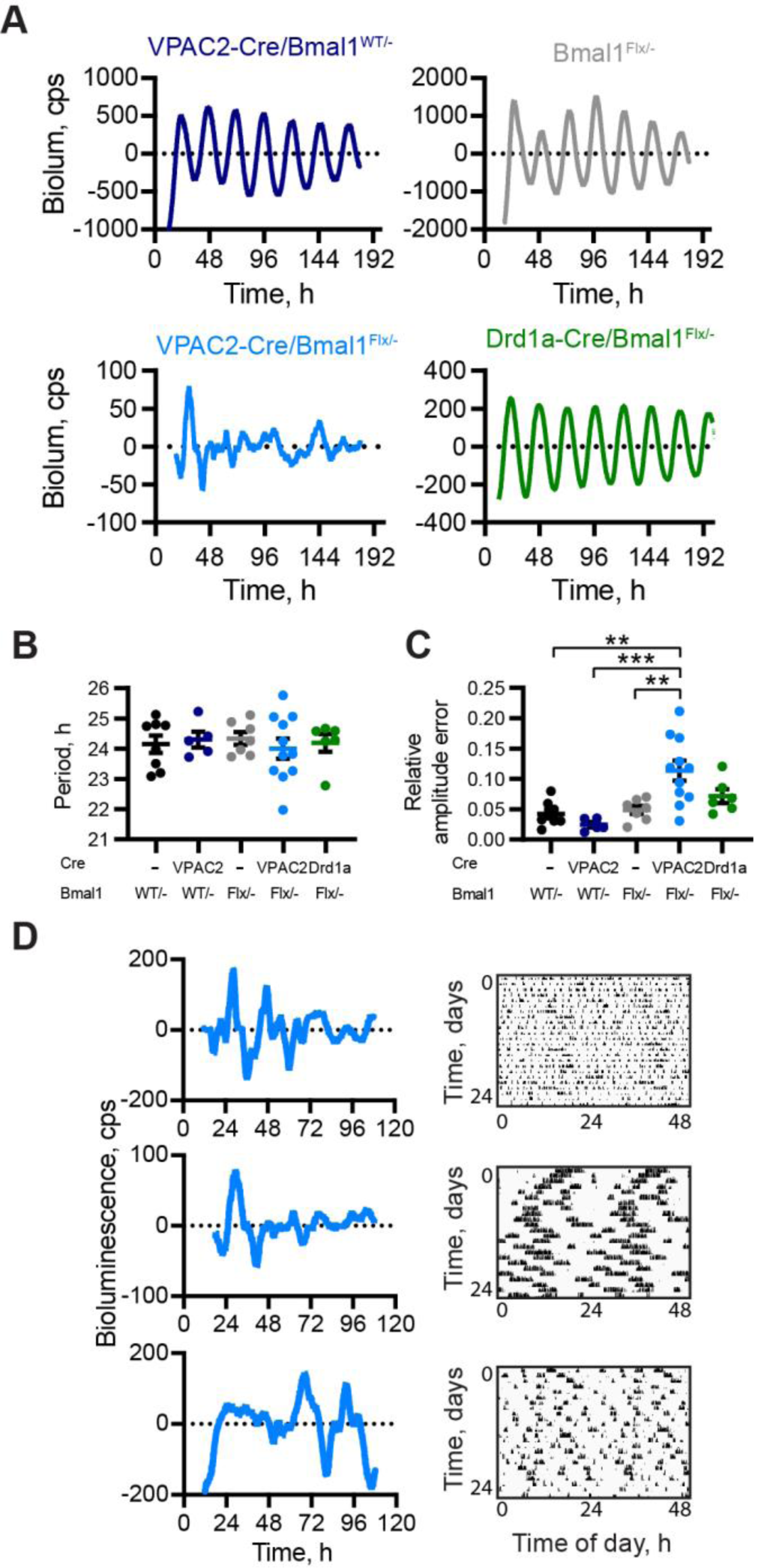
Deletion of BMAL1 from VPAC2-Cre-expressing cells compromises SCN molecular pacemaking. ***A***, Representative, baseline corrected PER2::LUCIFERASE bioluminescence traces from control, VPAC2-Cre/*Bmal1*^*flx/-*^, and Drd1a-Cre/*Bmal1*^*flx/-*^ SCN dissected in dim red light following recording of wheel-running rhythms in DD. ***B***, Period (mean ±SEM) of the first 4 bioluminescent cycles recorded from adult SCN slices; genotypes as in ***A***. n = 8 (*Bmal1*^*WT/-*^), 7 (*Bmal1*^*flx/-*^), 5 (VPAC2-Cre/*Bmal1*^*WT/-*^), 11 (VPAC2-Cre/*Bmal1*^*flx/-*^) and 6 (Drd1a-Cre/*Bmal1*^*flx/-*^). ***C***, Relative Amplitude Error (RAE) scores (mean ±SEM) for bioluminescent recordings from SCN slices. n as in ***B. D***, Representative PER2::LUCIFERASE bioluminescence rhythms from VPAC2-Cre/*Bmal1*^*flx/-*^ SCN slices alongside respective actograms from corresponding mice. All tests one-way ANOVA, with Tukey’s test for multiple comparisons. \**p* < 0.05, \*\**p* < 0.01, \*\*\**p* < 0.001.

## Discussion

The potency of the SCN as circadian pacemaker arises from network-level interactions that confer on it the emergent properties essential to its role. The SCN is, however, a heterogeneous mix of neuropeptidergic cell-types and so identifying the particular contribution(s) of defined cell populations to circuit-level function(s) is an important problem in circadian neurobiology. Here, we used intersectional genetics to manipulate the cell-autonomous clockwork of cells that express the VPAC2 receptor for VIP, and thereby demonstrated that these cells control circadian behaviour and molecular pacemaking within the SCN. These cells therefore constitute an important element of the network topology of the SCN, acting downstream of retinal and VIP-mediated signalling to maintain local and distant circadian coherence.

The ability of VPAC2 cells to determine behavioural circadian period complements results from other intersectional studies. Whereas lengthening the period of VIP cells (Vip-Clock^Δ19^) had no effect on free-running rhythmicity, the same manipulation in NMS cells (NMS-Clock^Δ19^), which include almost all VIP and AVP cells, lengthened behavioural period (Lee et al., 2015). Similarly, period-lengthening in Drd1a-Cre cells (including ∼60% of AVP cells) in Drd1a-Cre/*Ck1ε*^*Tau/Tau*^ mice concomitantly lengthened their behavioural period (Smyllie et al., 2016). Finally, directly altering the period of AVP cells alone accordingly shortened or lengthened behavioural rhythms (Mieda et al., 2016). Given that VPAC2 cells encompass ∼85% of AVP cells, the current period effects likely arise from altering cell-autonomous oscillators of VPAC2/AVP cells. Moreover, comparable effects can be seen from targetting AVP or VPAC2 cells in *ex vivo* preparations, insofar as the SCN period does not recapitulate behavioural periods. For example, *in vivo* period-lengthening of AVP cells to 26 h disappeared after the first cycle *ex vivo*, when SCN slices reverted to a wild-type 24 h. Nevertheless, dissection of core and shell SCN allowed the shell AVP cells to express their long cell-autonomous period, confirming effective genetic manipulation (Mieda et al., 2016). *Ex vivo*, SCN AVP cells are therefore less potent in period-setting than *in vivo*, similar to VPAC2-targetted SCN. Interestingly, however, VPAC2-Cre/*Ck1ε*^*Tau/-*^ slices showed a progressive lengthening of period, not seen in control slices, which happened without loss of precision or amplitude. This suggests that, with extended culture, the longer-period VPAC2 cells gained some influence over ensemble period, indicative of circuit-level plasticity.

Overall, therefore, the ability of VPAC2/AVP cells to determine circadian period is greatly enhanced in the intact animal compared to slices. While slice period variability may arise from technical factors such as Cre excision efficiency, the marked, consistent behavioural effect confirmed effective deletion. It is possible that VPAC2 cells function primarily as output cells of the SCN rather than dictating SCN periodicity per se, so their cell-intrinsic period is reflected at the behavioural level but not in slices. This would, however, require two separately oscillating populations within the SCN, which would be expected to lead to more unstable behavioural patterns than were observed. Instead, it is likely that modification of SCN function by behavioural state feed-back, at the level of the entire circuit or directly to the AVP and/or VPAC2 cells, underlies these differences. A precedent for behavioural feedback potentiating a compromised SCN is seen in arrhythmic VPAC2-null mice, which, after entrainment to a 24 h schedule of voluntary exercise, exhibit strong, free-running circadian behaviour on release into DD (Power et al., 2010). In the case of VPAC2-Cre/*Ck1ε*^*Tau/-*^ mice, it is likely that the sparse VPAC2 cells in the rest of the hypothalamus that receive input from the SCN, such as in the paraventricular nucleus (Kalló et al., 2004), would also have 24 h periods and may feed-back to, and resonate with, the intrinsically 24 h SCN VPAC2 cells. This feedback would reinforce the role of SCN VPAC2 cells in period-determination in vivo, leading to a change in behaviour, but would be absent in the slice.

In addition to their period-setting abilities, VPAC2 cells are also critical for rhythm maintenance, as evidenced by BMAL1 deletion in this cell population. Interestingly, the total number of targetted cells appears to be less important than the type of cell rendered circadian incompetent. In pan-neuronal deletion studies, loss of BMAL1 from ∼30% of neurons using a Nestin-Cre line did not have any effect on behavioural rhythms in DD (Mieda and Sakurai, 2011). Equally, deletion of BMAL1 in 65% of SCN cells using a single allele of Synaptotagmin10-Cre had no behavioural effect (Husse et al., 2011). Only in homozygous Synaptotagmin10-Cre mice, where BMAL1 was deleted from 83% of the SCN, was a behavioural phenotype evident. Moreover, in the current study loss of BMAL1 from Drd1a-expressing cells, which caused ∼70% loss of BMAL1 across the SCN, did not compromise circadian behaviour. Conversely, deletion of BMAL1 from NMS neurons, which constitute a minority of 40% of the SCN, nevertheless caused behavioural arrhythmia. In the current study, loss of BMAL1 from only 15-20% of total SCN cells, but 70% of VPAC2 cells, severely disrupted circadian behaviour. Unlike in pacemaking, where VPAC2 and AVP share many commonalities, loss of BMAL1 exclusively from AVP cells did not cause disrupted circadian behaviour, instead causing the nocturnal activity profile to widen and overall period to lengthen by ∼1 h (Mieda et al., 2015). Thus, VPAC2 cells appear to constitute a population in which rhythmicity is essential for overall circadian competence.

Further demonstration of the essential role of VPAC2 cells came from the *ex vivo* SCN slices, which exhibited highly disordered circadian cycles of TTFL activity, commensurate with the loss of behavioural coherence in these mice. This contrasts with the sustained TTFL function following the ablation of BMAL1 solely from AVP cells (Mieda et al., 2015), and from Drd1a-Cre cells in the current study. This suggests that the loss of behavioural coherence seen in *VPAC2-Cre/Bmal1*^*flx/-*^ mice has a more fundamental origin than simply an inability to convey behaviourally relevant circadian cues outside the SCN (a deficiency seen with loss of Prok2 receptor (Prosser et al., 2007). Even more strikingly, Ko et al. (2010) demonstrated that in global *Bmal1*^*-/-*^ slices, stochastic quasi-circadian rhythmicity can be seen, thus it is remarkable that loss of BMAL1 from 20% of the SCN can cause such a pronounced phenotype. This provides strong evidence that VPAC2 cells in the SCN exert powerful control over ensemble rhythmicity, a result complementing those of Patton et al. (2020), together showing that a competent TTFL in VPAC2 cells is necessary for coherent circadian oscillation.

Notwithstanding the overall effect of BMAL1 deletion, it is clear that the behavioural phenotype of VPAC2-Cre/*Bmal1*^*flx/-*^ mice was variable both in severity and the time that it took to be expressed under DD, suggesting that there are sub-populations of VPAC2 cells, as posited for other SCN populations (Kawamoto et al., 2003; Geoghegan and Carter, 2008). The advent of single cell transcriptomic profiling (Park et al., 2016; Wen et al., 2020) may reveal suitable markers for such sub-populations. That the behavioural disruption took time to emerge is intriguing, because, as with NMS-BMAL1 mice, phenotypes often appeared suddenly, with no obvious trigger, suggesting “*the presence of mechanisms capable of transiently compensating for the loss of molecular clocks*” (Lee et al., 2015). The initial rhythmicity *in vivo*, perhaps initiated by daily lighting cycles and/or sustained by light-driven behavioural rhythms, may have relied on coupling effects of non-VIP-ergic cells that compensated for the loss of cell-autonomous oscillators in the VPAC2 cells. Potential compensatory factors are AVP and GRP, both of which have been shown to be capable of inducing rhythmicity in VIP- or VPAC2-deficient SCN (Brown et al., 2005; Maywood et al., 2006, 2011). Levels of AVP, however, were reduced in *VPAC2-Cre/Bmal1*^*flx/-*^ mice, which may have exacerbated circadian disorganisation arising from loss of TTFL function in VPAC2 cells. The signalling molecules used by VPAC2 cells themselves remain unclear, not least because a global AVP knockout has only a minor effect on rhythmicity (Groblewski et al., 1981), although loss of AVP receptors does compromise intercellular coupling in the SCN (Yamaguchi et al., 2013). Similarly, NMS, another peptide marker for both core and shell SCN, is not essential for circadian function (Lee et al., 2015).

In summary, we have demonstrated that VPAC2 cells constitute an essential pace-setting and rhythm-generating population of the mammalian circadian system, necessary for fully coherent circadian oscillation both *in vivo* and *ex vivo*. Furthermore, we found that the strong pace-setting abilities of VPAC2 cells are disrupted, albeit not completely absent, on slice preparation, suggesting an unanticipated influence of extra-SCN populations in reinforcing periodicity. These findings complement those of Patton et al. (2020) that circadian competence in both VPAC2 cells and VIP cells is necessary for the *de novo* initiation of ensemble rhythms in SCN slices and behaviour. Thus, circadian competence in VPAC2 cells is *necessary* for rhythmicity, but they require their cognate signalling cell partners to be *sufficient* for rhythmicity. The VIPergic axis confers much of the fundamental robustness and intercellular communication that is essential to normal SCN function, and our work advances our knowledge of the individual functions of the cells inherent to this axis.

## Acknowledgements

The Authors thank Staff of the MRC LMB Biomedical Facility, Ares, for mouse breeding and handling, Staff of the Light Microscopy Facility for imaging support, and Emma Morris for experimental assistance. This work was funded by the Medical Research Council (MC_U105170643) core funding to MHH.

